# Transcriptome analysis reveals robust transcriptional reprogramming during early stages of *Cicer-Ascochyta* interaction

**DOI:** 10.1101/2022.03.26.485904

**Authors:** Ritu Singh, Aditi Dwivedi, Yeshveer Singh, Kamal Kumar, Aashish Ranjan, Praveen Kumar Verma

## Abstract

Ascochyta blight (AB) caused by a filamentous fungus *Ascochyta rabiei* is a major threat to global chickpea production. The mechanisms underlying chickpea response to *A. rabiei* remain elusive. Here, we investigated the comparative transcriptional dynamics of AB-resistant and susceptible chickpea genotypes upon *A. rabiei* infection to understand the early host defence response. Our findings revealed that AB-resistant plants underwent rapid and extensive transcriptional reprogramming compared to susceptible host. At early stage (24-hpi), mainly cell wall remodeling and secondary metabolite pathways were highly activated, while DEGs related with signaling components *viz.* protein kinases, transcription factors, and hormonal pathways show remarkable upsurge at 72-hpi, especially in resistant genotype. Notably, our data suggests imperative role of JA, ET, and ABA signaling in providing immunity against *A. rabiei*. Furthermore, gene co-expression networks and modules corroborated the importance of cell wall remodeling, signal transduction and phytohormone pathways. The hub genes such as MYB14, PRE6 and MADS-SOC1 discovered in these modules might be the master regulators governing chickpea immunity. Overall, we not only provide novel insights for comprehensive understanding of immune signaling components mediating AB resistance/susceptibility at early *Cicer-Ascochyta* interactions, but also offer a valuable resource for developing AB-resistant chickpea.

**Highlights:** Comparative transcriptomic and co-expression analysis of AB-resistant and susceptible chickpea genotypes reveals high-amplitude transcriptional dynamics in resistant plants, and also identifies TFs, PKs and phytohormone-crosstalk as core regulators for AB-resistance.

## INTRODUCTION

Plants are constantly confronted by various pathogens, and the outcome of this battle directly interferes with agricultural production. To combat pathogens, plants have evolved an array of sophisticated mechanisms consisting of constitutive and inducible immune systems (Jones and Dangl, 2006). On the other hand, phytopathogens employ diverse strategies for their survival, fitness, and to circumvent plant defence (Glazebrook, 2005; Toruño et al., 2016). Pathogen’s strategy varies with their lifestyle, where biotrophs need the host alive and necrotrophs want it to be dead. Evidences support the existence of a short biotrophic phase during necrotrophic infection as well (Chowdhury et al., 2017; Spanu 2012). The host immunity modulation by secreted effector proteins further challenges our understanding of plant responses to pathogen. Therefore, understanding the molecular basis of multifaceted plant-pathogen interactions remains prime focus to avert the crop loss.

The multi-layered plant immune system consists of preformed barriers, pathogen detection, and signal transduction components which coordinate the defence responses (Tsuda and Katagiri, 2010; Corwin and Kliebenstein, 2017). These pathogen-induced responses involve dynamic transcriptional reprogramming of multiple host processes leading to resistance or susceptibility. For instance, extensive degradation of cell wall polysaccharides is recognized by various receptor kinases which in turn trigger defence responses such as cell wall reinforcement at the pathogen penetration site by lignin and callose deposition (Brutus et al., 2010; Eynck et al., 2012; Flors et al., 2008). These responses activate reactive oxygen species (ROS) production, pathogenesis-related (PR) gene expression, and also the accumulation of plant defence hormones like salicylic acid (SA) and jasmonic acid (JA) (de Araujo et al., 2019). SA and JA are pivotal to combat biotic stress; however, emerging evidences indicate antagonistic or synergistic role of abscisic acid (ABA), ethylene (ET), Auxin (Aux) and brassinosteroid (BR) in immunity (Glazebrook, 2005; Pieterse et al., 2012; Catinot et al., 2015; Verma et al., 2016a).

Chickpea (*Cicer arietinum* L.) is one of the most widely grown legumes across the world, especially in arid and semi-arid climates (Singh et al., 2021a). A necrotrophic fungus, *Ascochyta rabiei* that causes Ascochyta blight (AB) disease, is a major constraint for global chickpea production. It infects all the aerial parts of the plant and causes up to 100% crop loss under conducive conditions (Knights and Hobson, 2004; Sharma and Ghosh, 2016). Further, the existence of teleomorph stage in *A. rabiei* accelerates their evolution, thus, actively defying the plant defence strategies (Mehmood et al., 2017). Consequently, *A. rabiei* isolates have developed resistance against quinone outside inhibitor (QoI) fungicides in Montana and North Dakota (Delgado et al., 2013; Owati et al., 2017). Importantly, resistance erosion has also been observed in several formerly known ‘AB-resistant’ chickpea cultivars (Sambasivam et al., 2020). This necessitates the identification of key genetic regulators of AB resistance in chickpea, which can be exploited for developing resistant plants. Previous expression studies provided limited understanding of chickpea response against *A. rabiei* infection (Coram and Pang, 2006; Jaiswal et al., 2011). An integrated transcriptome and degradome sequencing analysis uncovered numerous miRNAs and their targets that co-ordinate AB resistance in chickpea cultivars (Garg et al., 2019). However, the early-stage host transcriptional modulation that decides the fate of the battle during *Cicer-Ascochyta* interactions remains poorly understood. *A. rabiei* starts penetrating host tissue at 24-hours post-inoculation (hpi) followed by appearance of yellow spots on the leaves at 72-hpi (Ilarslan and Dolar, 2002). These stages are crucial to understand how AB resistant and susceptible plants respond to *A. rabiei* perception and penetration. Hence, in-depth investigations are required to appropriately comprehend the molecular components involved in resistance/susceptibility-related processes.

In this study, comparative transcriptome analysis of AB resistant (FLIP84-92C) and susceptible (PI359075) chickpea genotypes was performed at early stages (24- and 72-hpi) of *A. rabiei* infection. We identified differential gene expression pattern for defence-related components such as receptor kinases, phytohormone pathways, and transcription factors between both genotypes. In addition, the incorporation of weighted gene co-expression network analysis (WGCNA) revealed resistant/susceptible genotype-specific gene expression modules, and identified hub genes governing contrasting traits. Overall, we provide new insights into the complex *Cicer*-*Ascochyta* interactions, delineate AB-responsive host-physiological pathways, and deliver potential resistance genes for further characterization and downstream applications in chickpea improvement programs.

## MATERIALS AND METHODS

### Plant material, sample collection and RNA extraction

The seeds of FLIP84-92C(2) and PI359075 chickpea genotypes having contrasting phenotype for AB stress were obtained from Germplasm Resources Information Network (GRIN). FLIP84-92C is AB resistant *kabuli* chickpea variety with disease score of 2, whereas, PI359075 is a susceptible *desi* variety with 9 disease score (Tekeoglu et al., 2000). Plants were grown under control condition in growth chambers with 12 h light/dark cycle at 22 ± 2°C and >70% relative humidity. The *ArD2* isolate of *A. rabiei* (Indian Type Culture Collection No. 4638) was grown on chickpea-supplemented potato dextrose agar (PDA) plates. To isolate the fungal spores, *A. rabiei* grown plates were flooded with autoclaved distilled water, and surface was gently rubbed with a sterile loop. Obtained spore suspension was filtered through muslin cloth and spore concentration was determined using hemocytometer. For RNA-seq sample collection, the *A. rabiei* spore suspension containing 10^6^ spores/ml was sprayed on 14-day old chickpea plants. Plants sprayed with sterilized water were used as control. The aerial tissue of control and infected plants were collected at 24- and 72-hpi, immediately frozen in liquid nitrogen and stored at −80°C. Some of the infected plants from both the genotypes were monitored for two weeks for the progress of AB disease. Total RNA was isolated by utilising Plant RNeasy mini kit (Qiagen). The quantity and quality of the RNA samples were checked on 1% denaturing agarose gel, NanoDrop and Qubit Fluorometer. In total, 24 libraries (CFL, 24FL, 72FL, CPI, 24PI, 72PI, four replicates each) representing AB resistant (FLIP84-92C, denoted as FL) and susceptible (PI359075, denoted as PI) genotypes at control (C; uninfected) and AB stress (24- and 72-hpi; *A. rabiei-*infected) conditions, were sequenced on Illumina NextSeq500 platform using 2 x 150 bp chemistry.

### Data filtering and differentially expressed gene identification

The quality of raw reads was analyzed using FastQC (http://www.bioinformatics.babraham.ac.uk/projects/fastqc). Reads were quality-filtered and trimmed at the ends to remove low-quality bases and adapter sequences using FASTX-Toolkit (http://hannonlab.cshl.edu/fastx_toolkit). The reads having minimum PHRED score of 30 were considered for further analysis. The high-quality filtered reads were mapped on the reference genome of chickpea i.e. “CDC Frontier” (Varshney et al., 2013) using HISAT tool (Pertea et al., 2016) with default parameters. Further, the mapped reads were assembled via StringTie to generate the reference-guided assembly (Pertea et al., 2016). Read counts were computed using a python script prepDE.py3 (https://ccb.jhu.edu/software/stringtie/dl/prepDE.py3) which created count file containing the read count matrices for genes. Genes having less than 10 reads on an average per condition were excluded before the differential expression analysis. This filtered gene count matrix was used by a DESeq2 package (Love et al., 2014) in R for identification of differentially expressed genes (DEGs). DEGs were obtained for all pair-wise combinations (CPIvsCFL, CPIvs24PI, CFLvs24FL, CPIvs72PI, CFLvs72FL, 24PIvs24FL, 72PI vs72FL) with the criteria of adjP-value ≤ 0.05 and fold change (FC) ≥ 1.5.

### Functional enrichment and MapMan analysis

Annotation of the DEGs was performed using Blast2GO application of omics box (https://www.biobam.com/omicsbox). In order to identify the over-represented functional categories, gene ontology (GO) enrichment analysis was performed using PlantRegMap (Tian et al., 2020). Pathway enrichment analysis was performed by KOBAS web tool using hypergeometric model and significance threshold of *p*-value ≤ 0.05 (Xie et al., 2011). Venn diagrams, volcano plot and heat maps were prepared using TB Tool (Chen et al., 2020). Furthermore, MapMan analysis was conducted to visualize the *C. arietinum* gene expression data in the context of biotic stress pathways (Thimm et al., 2004). Mercator automated annotation pipeline (http://mapman.gabipd.org/web/guest/mercator) were used to assign *Cicer* genes to bins. The two-tailed Wilcoxon rank sum test adjusted by the Benjamin-Hochberg method (FDR ≤ 0.05) were utilized to define the differentially represented MapMan pathways.

### Family enrichment, clustering analysis and transcription factor binding-site prediction

Among DEGs, TFs, transcriptional regulators (TRs) and protein kinases (PKs) were identified using iTAK tool (Zheng et al., 2016). Fisher exact’s test in R (fisher.test function) was performed to identify significantly over-represented TF families in our datasets. The *p*-value ≤ 0.05 criterion was considered for significantly enriched families. The normalized count of differentially expressed transcription factors, regulators and protein kinases (DETFs, DETRs and DEPKs), respectively were used for hierarchical clustering analysis using R hclust package (version 3.6.2). Euclidean distance measure was used to calculate the distance between genes with ward.D as the agglomeration method. The cluster plots were generated with ggplot2 package in R (Wickham, 2016). Regulatory sequence analysis tools (RSAT)::Plants (http://plants.rsat.eu) was used to extract the promoter sequences of DEGs. The promoter sequences were subsequently submitted to binding site prediction tool available on PlantRegMap (http://plantregmap.gao-lab.org) to identify the TFs having maximum targets in the DEGs. Further, TF enrichment tool available on PlantRegMap website was used to find the TFs possessing significantly over-represented targets among the DEGs (*p*-value ≤ 0.05).

### Weighted gene co-expression network analysis

The co-expression network analysis was performed using the WGCNA package in R (Langfelder and Horvath, 2008). Normalized counts of genes in the upper 50% of coefficient of variation for expression across different conditions were used to construct the signed network. The degree distributions in each network followed the power law and satisfied the scale free topology criterion. The soft threshold power 34 was taken with a minimum module size of 30 genes, mergeCutHeight of 0.15 and deepSplit of 2. The WGCNA modules (co-expression networks) of eigengenes were identified as clusters of highly connected genes. Further, the networks correlated with traits were identified with the criterion of stability correlation *p*-value ≤0.05. The module significance cut off (Pearson’s correlation coefficient) ≥ 0.5 was considered, and the top most modules were retained for further analysis. Then, using the gene significance (GS) and module membership (MM) measurements, hub genes were identified with a cut-off GS and MM ≥ 0.8 and *p*-value ≤ 0.05 for a particular condition. Co-expressed genes having weight ≥ 0.2 were visualised using Cytoscape v3.8.2. The top genes identified through cytoHubba and network analyser were highlighted in the regulatory network.

### qRT-PCR analysis

To validate the result of RNA-seq we performed quantitative real time-polymerase chain reaction (qRT-PCR) analysis of 12 randomly selected genes. The gene specific primers were designed through Primer Express software (Applied Biosystems, USA). The qRT-PCR was carried out on 7900HT Sequence Detection System (Applied Biosystems, USA) using SYBR Green and 50°C for 2 min, 95°C for 10 min, 40 cycles of 95°C for 15 s, and 60°C for 1 min cycling conditions. The gene expression values are derived from three biological replicates having three technical replicates. Chickpea β-tubulin and EF-1α gene were used as an internal control to normalise the data.

## RESULTS

### Transcript profiling of AB resistant and susceptible chickpea genotypes at early stages of *Ascochyta* infection

To understand the host transcriptional dynamics upon *A. rabiei* infection, transcriptome profiling of resistant (FLIP84-92C) and susceptible (PI359075) chickpea genotypes under control and AB-stress conditions was performed at 24- and 72-hpi. The details of raw, filtered and mapped reads on the reference genome are given in Dataset S1. To identify the DEGs, seven pair-wise comparisons were performed between control, 24- and 72-hpi time-points and genotypes. Four sets represent transcriptional reprogramming within the genotype in response to *A. rabiei* [CPIvs24PI, CFLvs24FL, CPIvs72PI, CFLvs72FL; samples are designated as ‘condition genotype’ (CPI)]. Three sets show differential response between the susceptible and resistant genotypes (CPIvsCFL, 24PIvs24FL, 72PIvs72FL). Overall fold change distribution of identified genes is shown in Figure S1.

In response to AB-stress, a total of 332 (177 up- and 155 down-regulated) and 980 (625 up- and 355 down-regulated) genes were differentially expressed in susceptible plants at 24- and 72-hpi, respectively. However, in resistant genotype, 441 (203 up- and 238 down-regulated) and 8908 (4602 up- and 4306 down-regulated) genes showed differential expression (Figure 1A). Both genotypes showed more DEGs at 72-hpi, though the number was much higher in resistant genotype suggesting rapid and dramatic transcriptional changes. Although, 3.88% (24-hpi) and 5.73% (72-hpi) AB stress-specific DEGs were shared between the resistant and susceptible genotype, their expression levels were not correlated (r<0.4). Among them, 39 showed opposite expression pattern in both the genotypes (Figure 1C). In contrast, a total of 93.1% and 93.6% DEGs were uniquely expressed in resistant genotype, while 90.9% and 42.1% DEGs were exclusive to susceptible genotype at 24- and 72-hpi, respectively. Further, the shared DEGs between 24- and 72-hpi in resistant (3.21%) and susceptible (10.53%) genotypes represent the constitutively activated defence mechanisms during various infection stages (Figure 1B).

**Figure 1:**
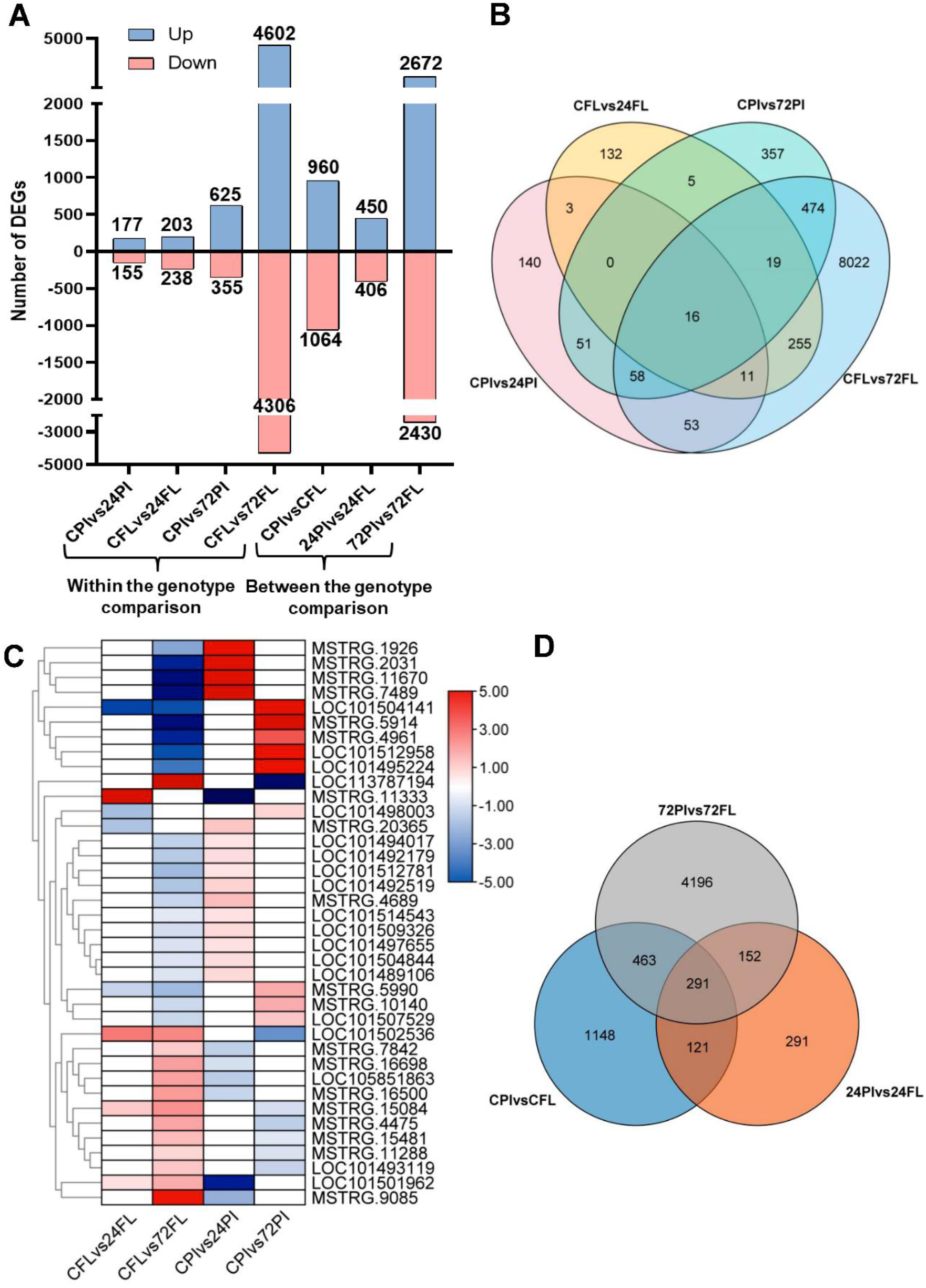
Transcriptome profile of resistant (FLIP 84-92C) and susceptible (PI 359075) chickpea genotypes in response to *Ascochyta rabiei* infection. **A,** Number of differentially expressed genes (DEGs) identified within the genotype (CPIvs24PI, CFLvs24FL, CPIvs72PI, CFLvs72FL) and between the genotype comparisons (CPIvsCFL, 24PIvs24FL, 72PIvs72FL) at 24 and 72 hpi under control and AB stress condition. **B,** Venn diagram depicting overlapping and specific DEGs within genotypes at both time-points. **C,** Heatmap showing expression of shared genes having opposite expression pattern in resistant and susceptible genotype. **D,** Venn diagram depicting overlapping and specific DEGs between the genotypes.

Similarly, when resistant and susceptible genotypes were compared, the number of DEGs varied from 856 (450 up- and 406 down-regulated; 24-hpi) to 5102 (2672 up- and 2430 down-regulated; 72-hpi) (Figure 1A). Interestingly, few genes (10.9%) overlapped between the control and AB-stress condition, thus showing a major proportion of AB-induced DEGs. Moreover, 4.4% and 63% DEGs were exclusive to 24- and 72-hpi, respectively (Figure 1D). DEGs with specific induction or suppression upon infection could mediate the variable defence responses exhibited by these contrasting genotypes. Overall, results illustrated robust transcriptional reprogramming at early stages of infection in chickpea; however, the host response differs between incompatible (resistant) and compatible (susceptible) interactions.

### Differential transcriptional modulation of host defence pathways upon pathogen attack

To gain insights into *A. rabiei*-triggered pathways in these chickpea genotypes, functional classification and pathway assignment of DEGs were performed. In both genotypes, the up-regulated DEGs were mostly associated with defence, response to biotic stimulus, hormone signaling, oxidation-reduction, protein phosphorylation and modification (Dataset S2). However, DEGs associated to each GO category were higher in resistant genotype suggesting robust activation of defence components. Up-regulated genes in FLIP84-92C were mainly associated with defence response by cell wall thickening, callose deposition, and regulation of fatty acid oxidation, while developmental and reproductive process were down-regulated. In contrast, up-regulated genes in PI359075 genotype were mainly involved in toxin, phytoalexin, camalexin and flavonoid biosynthetic processes. Notably, when resistant genotype was compared to susceptible; ‘photosynthesis’, photosystem II assembly’, ‘photosynthetic electron transport chain’ and ‘photosystem II oxygen evolving complex assembly’ processes showed up-regulation in resistant plants, indicating differential regulation of photosynthetic processes between both the genotypes. Moreover, enrichment of resistant genotype-specific DEGs suggests their association with translation, ribosome biogenesis, and lipid biosynthesis processes, however, susceptible genotype-specific DEGs belong to cellular carbohydrate metabolic, small molecule and vitamin biosynthetic process (Figure S2). Such variation in gene regulation upon *Ascochyta* infection reflects the different strategies tailored by these genotypes.

Further, categorization of DEGs into MapMan biotic stress pathway revealed minimal transcriptional alterations at 24-hpi in both the genotypes compared to 72-hpi, though relatively large number of genes was differentially regulated in resistant genotype (Figure S3). When resistant genotypes were compared to susceptible, considerable up-regulation of redox state, glutathione-S-transferase (GST), signaling, secondary metabolites, PR proteins and proteolysis pathways were observed at 24-hpi. Consistently, signaling and proteolysis pathways were differentially regulated at 72-hpi, but showed slight changes in susceptible genotype (Figure 2A-B). Peroxidases and β-glucanase were mostly down-regulated. MAPKs along with WRKY and MYB TFs were also up-regulated in both the genotypes, although the number of genes is substantially higher in resistant plants. ERF and DOF TFs predominantly down-regulated but bZIP TFs were induced in resistant plants, whereas, no obvious changes were observed in susceptible genotype. In addition, genes associated to SA, JA, ET, and ABA were up-regulated in resistant genotype, while BR and Aux-related genes mainly showed down-regulation. On the other hand, only few hormone-related genes were differentially expressed in susceptible genotype. These observations are concordant with the enrichment analysis (Dataset S2).

**Figure 2:**
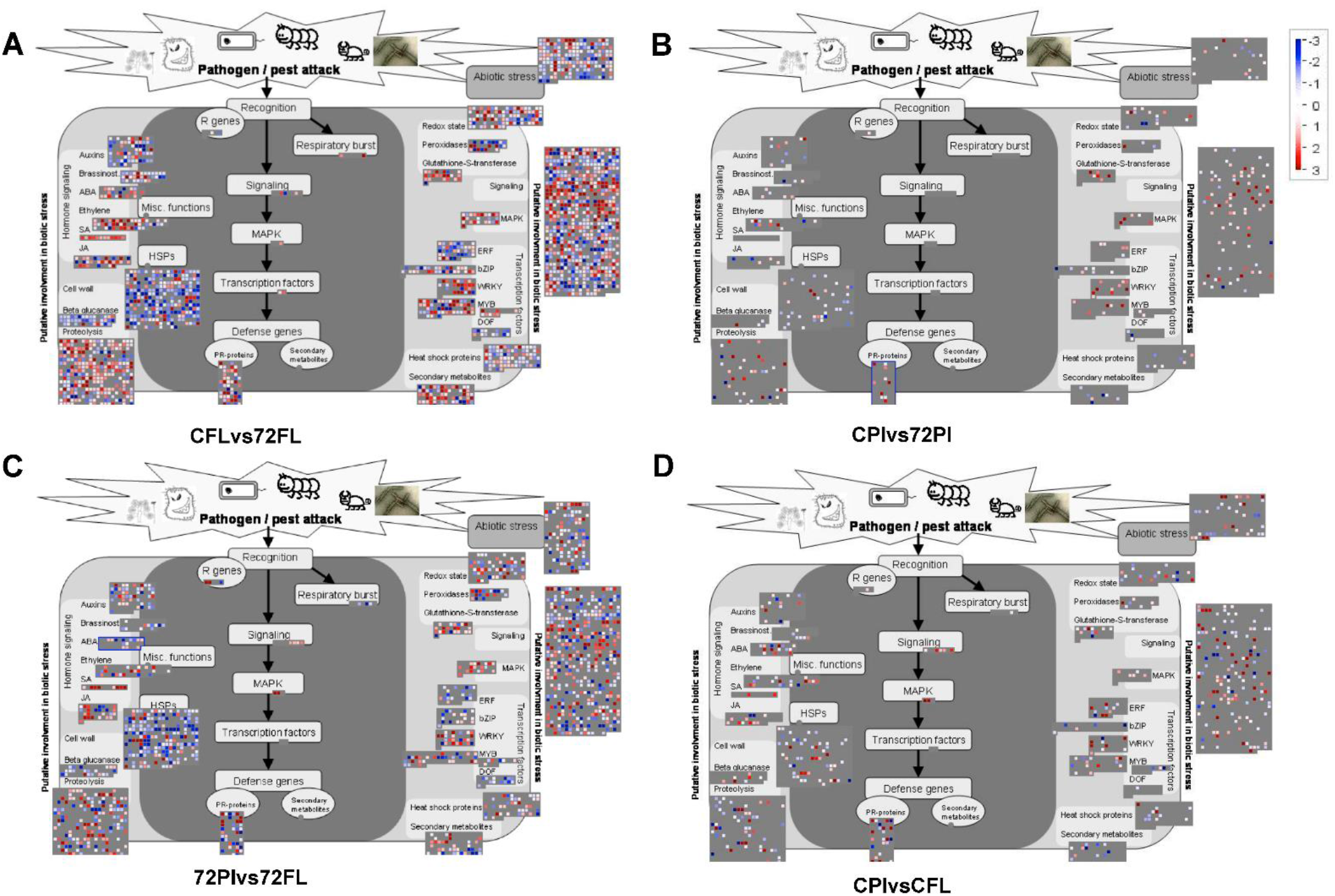
Overview of biotic stress pathways of DEGs visualized by MapMan. Each square represents an individual gene. Upregulated and downregulated genes are represented in red and blue, respectively. The scale bar shows fold change values. **A and B,** show altered pathways in resistant (CFLvs72FL) and susceptible (CPIvs72PI) genotype at 72 hpi of *A. rabiei*. **C,** CPIvsCFL shows altered pathways in resistant genotype compared to susceptible under control condition. **D,** 72PIvs72FL shows differentially regulated pathways between resistant and susceptible genotypes at 72 hpi.

Moreover, various abiotic stress-related DEGs were also identified, which could be common mediators of biotic and abiotic stress responses. Intriguingly, resistant genotype showed similar differentially regulated pathways in both comparisons i.e. CFLvs72FL and 72PIvs72FL (Figure 2A and 2C), indicating minimal response of susceptible genotype against *A. rabiei*. Differential expression of JA, ET, and ABA-related genes along with WRKY, ERF, and MYB TFs was observed in genotype-level comparisons under control condition, implying pre-activated basal defence mechanisms in resistant genotype (Figure 2D). Further, these RNA-seq data were validated by qRT-PCR analysis (Figure S4, Dataset S3). Overall, these analyses demonstrate robust activation of diverse defence mechanisms in resistant genotype, and highlight the importance of signal transduction and gene regulatory machinery co-ordinating high-amplitude transcriptional reprogramming against pathogen.

### Protein kinases and Transcription factors are extensively implicated in host defence

PKs and TFs are important players in perception and transduction of stress signals for coordinated defence. In our data, 520 DEPKs, and 905 DETFs/DETRs were identified, which were most abundant in resistant genotype at 72-hpi (Figure 3A; Dataset S4). Among them 35 DETF/DETR and 63 DEPK families/sub-families were uniquely expressed in resistant genotype upon AB-stress (Figure S5A and S6A). Between genotype comparisons showed 17 unique AB stress-specific DETF and DETR families along with 37 DEPKs in the resistant genotype (Figure S5B and S6B). These families include LIM, MADS-M type, C2C2-LSD, C2C2-CO-like, E2F-DB and DBB TFs along with LUG, SWI/SNF-SW13 and Pseudo-ARR-B TRs. PKs include receptor like kinase (RLK)-Pelle_LRR sub-families V, IV, Xb1 and XIIIa, CMGC_GSK and TTK.

**Figure 3:**
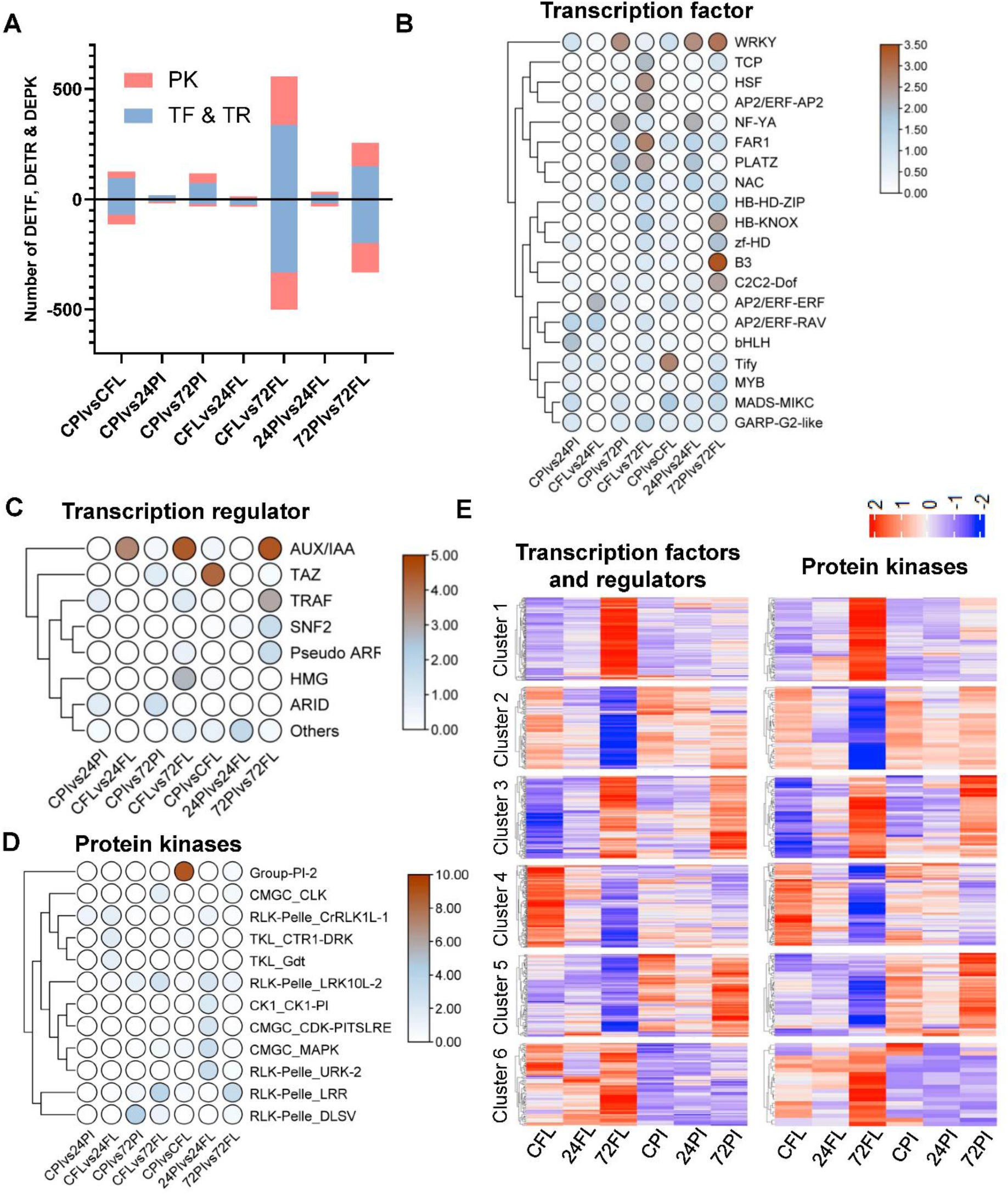
Differentially expressed transcription factors (DETFs), transcription regulators (DETRs) and protein kinases (DEPKs) under AB stress. **A,** Total number of TFs, TRs and PKs in DEGs. **B, C and D,** represent the heatmaps of enriched TF, TR and PK families, respectively. **E,** Clustering of DETFs, DETRs and DEPKs under control and AB stress conditions. The clustering was performed on DeSeq2 normalized gene count matrix under different combinations. Genes showing similar expression trend were grouped together into six clusters (Cluster 1-6). Cluster 1 and 2 show DETFs, DETRs and DEPKs at 72 hpi in resistant genotype (FLIP 84-92C). Cluster 3 represents 72 hpi specific cluster in resistant and susceptible (FLIP 84-92C and PI 359075) genotypes. Cluster 4 depicts differential expression trend between control and 72 hpi in resistant genotype. Cluster 5 and 6 are susceptible and resistant genotype specific cluster, respectively.

Further analysis showed significant enrichment of 20 TF, 8 TR and 12 PK families at least in one of the datasets (Figure 3B-D). Majorly TRs (Aux/IAA, TRAF, SNF2, Pseudo-ARR-B, HMG) and PKs (RLK-Pelle_LRR, CMGC_MAPK, CMGC_CDK) were enriched in resistant genotype. WRKY, NAC, PLAZ, and FAR1 TFs were enriched in both the genotypes under AB-stress, whereas MYB, B3, TCP, AP2/ERFs, HSF, GARP-G2, MADS-MIKC, B3, C2C2-DOF, HD-HD-ZIP and zf-HD families were exclusively enriched in resistant genotype. Moreover, hierarchical clustering identified six expression trend clusters of DETFs, DETRs and DEPKs (Figure 3E). Cluster 1 shows up-regulated expression in resistant genotype at 72-hpi which mainly contains WRKY and B3 TFs and DLSV, LRR, RLCK and WAK PKs. Lr10 was the most dominant member in this cluster. Resistant genotype-specific cluster 2 includes down-regulated Aux/IAA, bHLH, MYB, HB-HD-ZIP TFs and RLK-Pelle_LRR PKs. Cluster 3 is specific to 72-hpi showing induced expression in both the genotypes. This cluster harbors bHLH and NAC TFs along with RLK-Pelle_DLSV PKs. Interestingly, cluster 4 demonstrates down-regulation at 72-hpi, while up-regulation under control conditions in resistant genotype (Figure 3E). In this cluster, bHLH, GATA and AP2/ERF-ERF TFs along with RLK-Pelle_LRR and serine/threonine kinase were present. Cluster 5 and 6 represent susceptible and resistant genotype-specific up-regulation, respectively. In cluster 5, mainly bHLH, AP2/ERF-ERF, C2H2 TFs and probable inactive receptor kinases were identified, whereas cluster 6 contains large number of WRKY TFs and RLK-Pelle PKs (Dataset S5). Noticeably, most of TFs and PKs were exclusive to one genotype, while others showed similar expression pattern in both genotypes, implicating both common and specific response to *Ascochyta* infection in resistant and susceptible host. Data also suggests that DEPKs and DETFs between both the genotypes might be the main driving force for the differential expression of other defence-related genes.

The gene expression dynamics is tightly controlled by TFs which bind to specific *cis*-regulatory regions of target genes. In our data, targets of MADS, ERF, MYB, NAC, and WRKY TFs were over-represented upon infection, suggesting their role in plant immunity (Dataset S6). MYB14 and MYB36 TFs enriched in our datasets target phytohormone related genes, thus suggesting intricate role of hormones during *Cicer-Ascochyta* interactions. Along with pathogenesis-related gene transcription activator (*PTI6*), targets for MADS-box protein *SUPPRESOR OF OVEREXPRESSION OF CONSTANS1* (*SOC1*) and *BASIC-PENTACYSTEINE* (*BPC*) were markedly enriched among the DEGs. The *SOC1* and *BPC* regulate flower and leaf development-related genes in arabidopsis (Lee et al., 2008; Shanks et al., 2018). However, defence-related targets of these TFs including disease resistance protein RPS5, thaumatin-like protein PR5a, PR10, WRKY TFs along with hormone-related genes were also abundant in our datasets, indicating their potential role in plant immunity which remains to be explored. Interestingly, growth and development pathways were down-regulated in resistant genotype upon *A. rabiei* exposure (Dataset S2). Thus, these genes might be crucial players for growth-immunity trade-off.

### Modulation of hormonal pathway genes highlights central role of JA, ET and ABA signaling in response to blight fungus

The phytohormone cross-talk plays significant role in defence against invading pathogens. In our datasets, we identified 7, 33, 34, 41, 54 and 90 DEGs associated with SA, ABA, BR, JA, ET and Aux pathways, respectively (Dataset S7). JA biosynthesis genes *viz.* lipoxygenase (LOX), allene oxide synthase (AOS), allene oxide cyclase (AOC) and 12-oxophytodienoate reductase (OPR) were differentially expressed in resistant genotype (Figure 4A). LOX and OPRs were up-regulated, whereas, AOS and AOC were down-regulated upon *A. rabiei* infection in resistant genotype. Moreover, 1-aminocyclopropane-1-carboxylate (ACC) synthase and ACC oxidase were up-regulated suggesting induced biosynthesis of ET (Figure 4B). In ABA biosynthesis, zeaxanthin epoxidase (ZEP) was down-regulated, while short-chain alcohol dehydrogenase/reductase (SDR) and abscisic aldehyde oxidase (AAO) were induced in both the genotypes (Figure 4C). SA, BR and Aux biosynthesis genes were not differentially expressed. Furthermore, ABA-8’-hydroxylase was down-regulated in both the genotypes, while jasmonate methyltransferase (JMT) and salicylate carboxymethyltransferase (SAMT) were exclusively induced in resistant genotype at 72-hpi (Figure 4D). This suggests the activated biosynthesis of methyl jasmonate (MeJA) and SA-methyl ester (MeSA) for JA- and SA-mediated signaling during plant defence.

**Figure 4:**
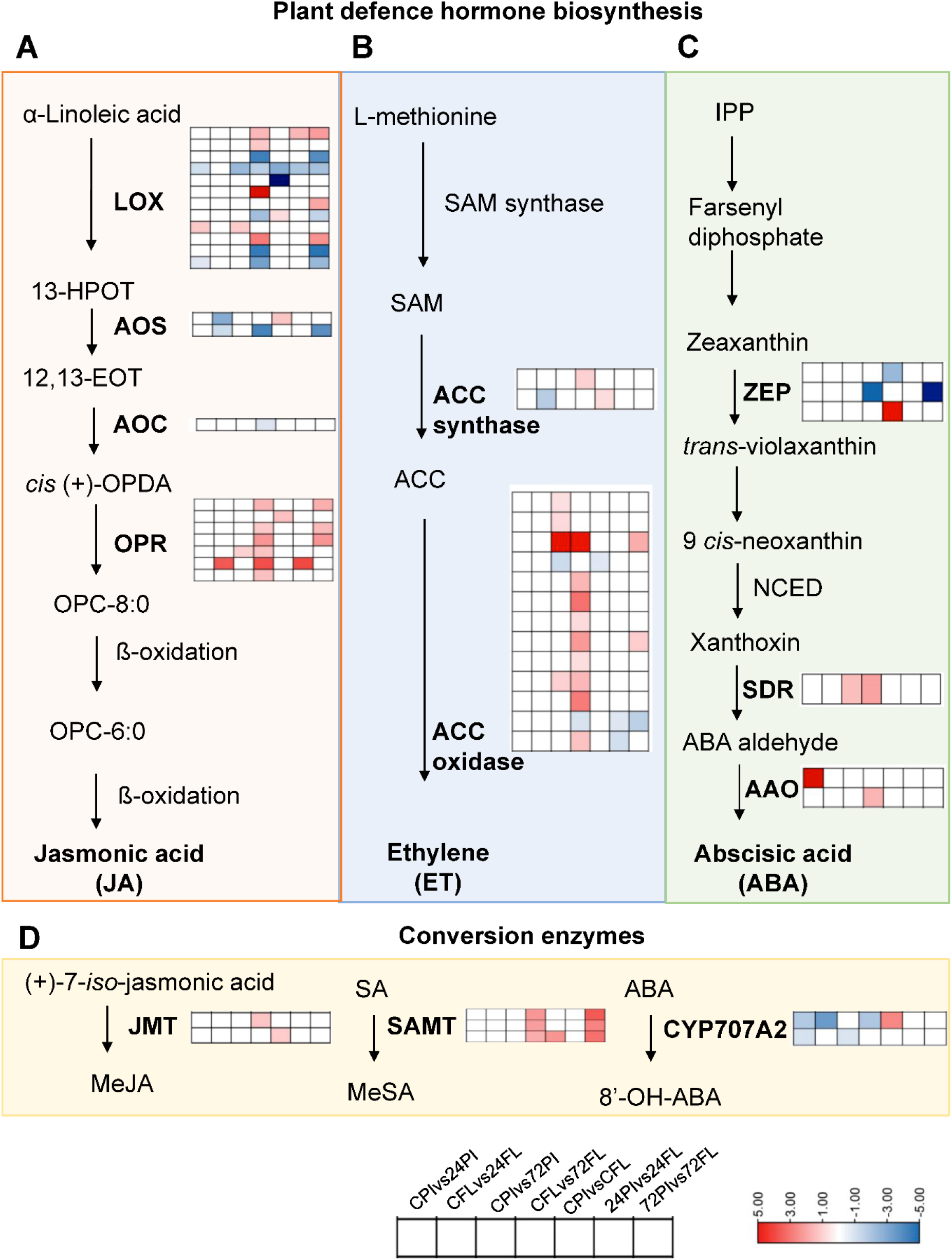
Modulation of plant defence hormones in response to AB stress. **A, B and C,** illustrate the jasmonic acid (JA), ethylene (ET) and abscisic acid (ABA) biosynthesis pathways, respectively, along with the heatmap of the genes showing alteration during AB stress. The scale represents the log_2_ fold change for comparison datasets. **D,** represents the enzymes detected in conversion pathways of hormones. LOX- Lipoxygenase; 13-HPOT-(13S)- hydroperoxyoctadecatrienoic acid; AOS- allene oxide synthase; 12,13- EOT- 12,13-epoxy- octadecatrienoic acid; AOC- allene oxide cyclase; cis (+)-OPDA- cis-(+)-12-oxophytodienoic acid (OPDA); OPR- OPDA reductase; OPC- 8:0-12-oxophytoenoic acid; SAM- S-adenosyl- methionine; ACC- 1-aminocyclopropane-1-carboxylic acid; ACS-ACC synthase; ACO- ACC oxidase; IPP- isopentenyl diphosphate; ZEP- zeaxanthin epoxidase; NCED- 9-cis-epoxy carotenoid dioxygenase; SDR- short-chain alcohol dehydrogenase/reductase; AAO- abscisic aldehyde oxidase; JMT- jasmonic acid carboxyl methyltransferase; MeJA- Methyl jasmonate; SA- salicylic acid; MeSA- Methyl salicylate; SAMT- S-adenosyl-l-methioninecarboxyl methyltransferase; CYP70742- ABA-8′-hydroxylases; 8′-OH-ABA- 8′-hydroxy ABA.

Moreover, DEGs associated with JA, SA, ABA and ET regulatory pathways were up-regulated, whereas, Aux and BR were down-regulated in resistant genotype at 72-hpi (Figure S7; Dataset S7). At 24-hpi fewer changes were observed. SA response-associated DEGs; methylxanthosine synthase, pentatricopeptide repeat-containing protein and glycosyltransferase were exclusively up-regulated at 72-hpi in resistant genotype. Intriguingly, many DEGs were shared in SA and JA pathway which is suggestive of possible cross-talk. In resistant genotype, ethylene response factors (ERFs) were differentially expressed, whereas CBS-domain containing protein, ethylene receptor, and codine-O-demethylase clearly over-expressed. In addition, ABA-responsive genes *viz.* GEM protein, HVA22, serine/threonine phosphatases were also up-regulated, whereas, auxin-responsive (SAUR and Aux/IAA) and BR-related genes were down-regulated in resistant plants. Further, auxin transporters were differentially regulated indicating the altered auxin levels during fungal infection. The elevated auxin levels are correlated with Fusarium Head Blight (FHB) susceptibility in wheat (Wang et al., 2018).

These observations were also supported by the GO enrichment analysis. In susceptible genotype, ‘response to JA’ and ‘response to SA’ terms were over-represented among up-regulated DEGs, while ‘response to JA’, ‘response to ABA’, ‘SA-mediated signaling pathway’, ‘ethylene-activated signaling pathway’, and ‘response to ethylene’ were enriched in resistant genotype. Among the hormone-responsive terms mainly WRKY40, MYB33, MYB48, ARR2 TFs, PR-proteins, class I and V chitinases were identified in resistant genotype. On the other hand, WRKY43, NAC05, and MYB48 TFs were present in susceptible genotype. Down-regulated terms include ‘auxin-activated signaling pathway’ and ‘auxin polar transport’ in resistant genotype, whereas, ‘response to brassinosteroid’ was commonly represented in both the genotypes (Dataset S2). Among the auxin and brassinosteroid response terms only AP2/ERF-AIL5 and NAC32 TFs were present, respectively. Identification of these TFs and defence genes under hormone response category implicates their role in downstream activation of host processes. Collectively, our findings demonstrate induced biosynthesis and signaling of JA, ET, and ABA in resistant genotype. The SA, Aux and BR biosynthesis genes remain unchanged. Moreover, the regulatory genes of SA showed up-regulation, while, Aux and BR were down-regulated during *A. rabiei* infection. This implicates that JA, ET, ABA, and their plausible cross-talk, are central regulators of chickpea defence against AB-stress. Further, these phytohormones modulate various genes through signaling cascades which is indicative of complex co-expression programs that work together to defend the host plant.

### AB resistance and susceptibility associated co-expression modules in chickpea

Further, we performed WGCNA analysis to divulge the modules and gene regulatory networks (GRNs) associated with particular genotype and time-points. Correlation of module eigengene with sample trait (i.e. resistance and susceptibility) identified 12 co-expression modules (comprised of 39 to 5425 genes) associated with particular condition (Figure 5A). For further analysis, highly correlated (r≥0.5 and ≤-0.5) and significant modules (*p*-value≤0.05) were selected in each dataset. Under control condition, the strongest positive correlation (r=0.91) was obtained for the lightgreen module in resistant genotype. Functional categorization of this module showed over-representation of ‘defence response to fungus’, ‘response to biotic stimulus’, ‘plant-pathogen interaction’ and ‘MAPK signaling pathways’, signifying the basal pre-formed defence in resistant genotype (Figure 5B-C).

**Figure 5:**
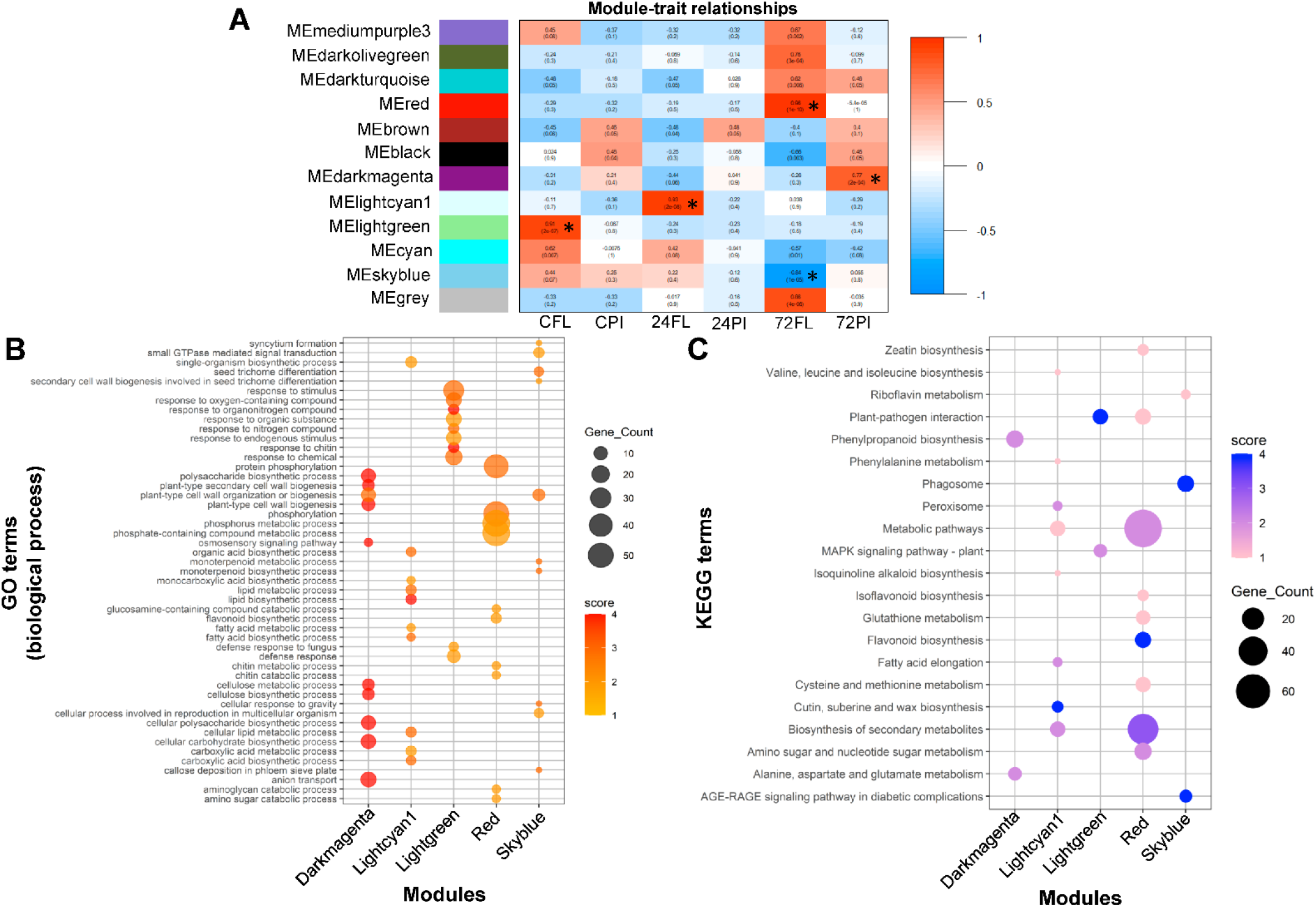
Gene co-expression module identification and functional category enrichment analysis. **A,** The correlation matrix of module eigengene values and phenotypes; red and blue color denotes positive and negative correlation with gene expression, respectively. **B,** GO enrichment analysis showing top ten biological process terms enriched in at least one module. **C,** KEGG pathway analysis of the genes identified under various significant modules.

Under AB stress, lightcyan1 (r=0.93) and red (r=0.96) modules were identified at 24- and 72-hpi, respectively, showing strong positive correlation with resistance trait. Higher number of genes was identified in red module (638) compared to lightcyan1 (39), which is concordant with our differential gene expression data. Lightcyan1 module over-represents monocarboxylic acid, lipid and fatty acid biosynthetic, fatty acid elongation, cutin, suberin, wax, isoquinoline alkaloid and secondary metabolites biosynthesis-related terms, suggesting rapid cell wall remodeling and secondary metabolite production in resistant chickpea to restrict fungal penetration. At later stage (red module), ‘response to wounding’, ‘flavonoid biosynthetic and metabolic process’, ‘chitin metabolic process’ ‘protein phosphorylation’, ‘biosynthesis of secondary metabolites’, ‘glutathione metabolism’, ‘zeatin biosynthesis’ and ‘plant-pathogen interaction’ terms were over-represented. Moreover, a negatively correlated skyblue module (r=-0.84) was identified at 72-hpi, harboring gene-set belonging to ‘callose deposition in phloem sieve plate’, ‘cell wall organization or biogenesis’, and ‘monoterpenoid metabolic and biosynthetic process’ categories in resistant genotype (Figure 5B-C). The data indicates extensive involvement of post-translational modifications via kinase activity, antioxidant metabolism and flavonoid biosynthesis pathways in providing resistance to AB disease, which is in agreement with our DEGs data.

In susceptible genotype, a positively correlated (r=0.77) darkmagenta module was detected at 72-hpi showing over-representation of ‘polysaccharide biosynthetic process’ ‘plant-type secondary cell wall biogenesis’, ‘alanine, aspartate, and glutamate metabolism’ and ‘phenylpropanoid biosynthesis’ (Figure 5; Dataset S8). Interestingly, ‘protein phosphorylation’ category was also enriched; however, the number of genes was much higher in resistant (red; 50) than susceptible genotype (darkmagenta; 32). This signifies the role of PKs in regulating the function of downstream genes mediating AB resistance, which is consistent with DEPK analysis. Noticeably, no significant modules were identified under the control and 24-hpi conditions in susceptible genotype, which further indicates the delayed or suppressed defence (Figure 5A). Overall, enrichment analysis presents an overview of the reprogrammed chickpea defence pathways, and indicates the presence of genotype-specific hub genes mediating differential response of both genotypes.

### Core regulators orchestrating the host response against AB disease

To identify the core regulators governing AB resistance/susceptibility, we performed GRN and GS-MM analysis (Figure 6). The highly connected hub genes in GRN could be master regulators of the module and hence, the associated trait. GS-MM analysis identified trait-associated hub genes from lightgreen (66), lightcyan1 (25), red (356), skyblue (187), darkmagenta (53) modules (Figure S8; Dataset S9). In accordance to our earlier observations, the highly connected red GRN showed remarkable prevalence of defence-related transcripts and signaling components among hub genes, indicating complex regulatory mechanisms to protect the plant from pathogen (Dataset S10). Moreover, extensive functional category enrichment of hub genes provided crucial insights into the differential response regulators in AB-resistant and susceptible genotypes.

**Figure 6:**
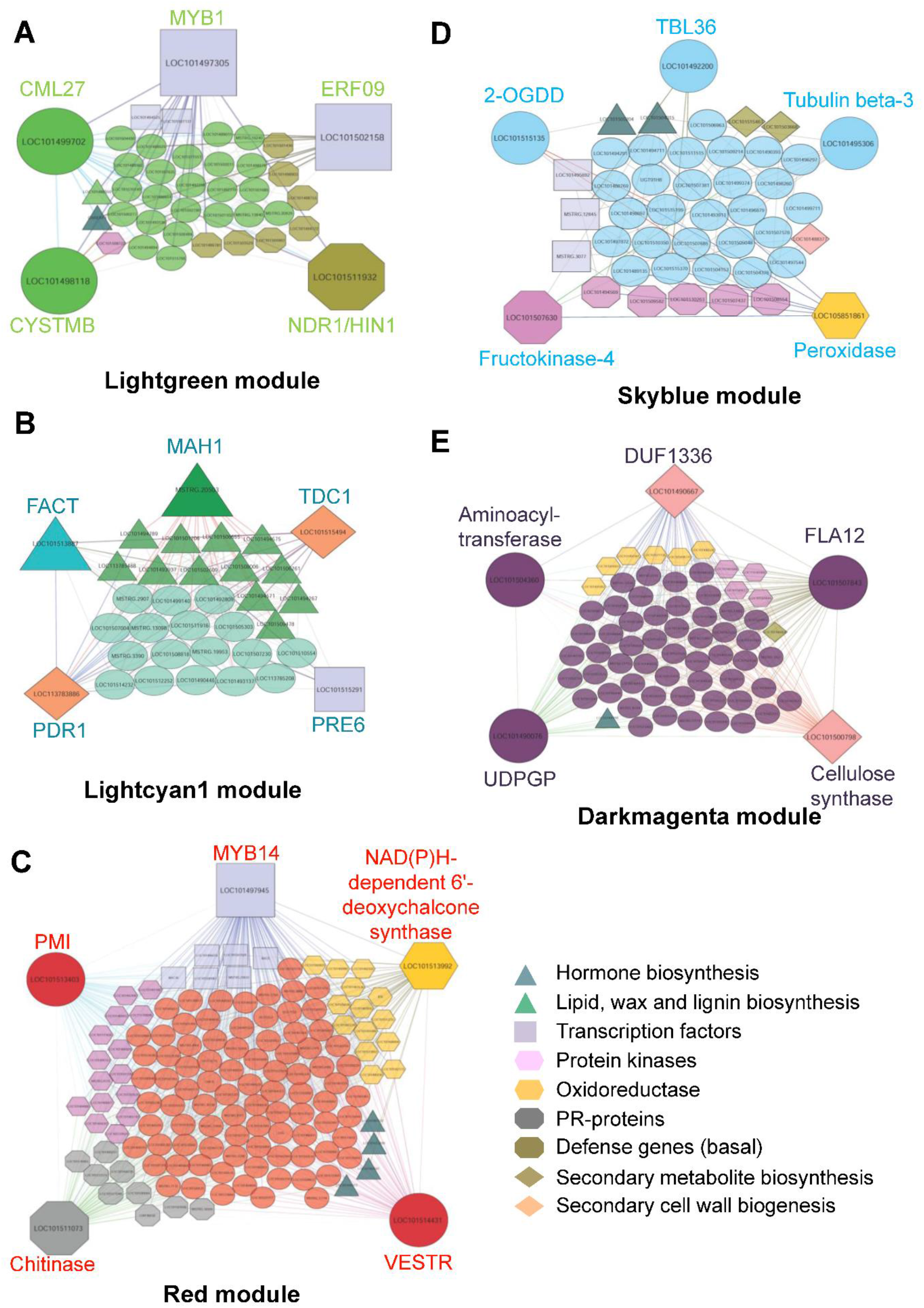
Transcriptional regulatory networks operating in resistant and susceptible genotype under AB stress. **A-E,** The regulatory network of genes having weight ≥ 0.2 are shown and top five hub genes are highlighted by bigger shapes in each module. Edges represent known interactions between the genes. CML27- calcium-binding protein CML27; CYSTMB- cysteine-rich and transmembrane domain-containing protein B; NDR1/HIN1- NDR1/HIN1-like protein 13; MAH1- alkane hydroxylase MAH1; FACT- fatty alcohol:caffeoyl-CoA acyltransferase; PDR1- pleiotropic drug resistance protein 1; TDC1- tyrosine decarboxylase 1; PMI- Plastid movement impaired protein; VESTR- vestitone reductase; 2-OGDD- 2- oxoglutarate-dependent dioxygenase; TBL 36- protein trichome birefringence 36; FLA12- fasciclin-like arabinogalactan protein 12; UDPGP- UTP-glucose-1-phosphate uridylyltransferase.

### Basal defence is pre-activated in AB-resistant genotype

Basal expression of defence genes has an imperative role in preventing fungal invasion. Notably, no significant module was detected in susceptible genotype; however, the hub genes identified under resistant (lightgreen) module were mainly involved in defence response, immune system process, and plant-pathogen interaction. These genes include 3-ketoacyl-CoA synthase, ERF027 and ABA-8’-hydroxylase. Genotype-level differences under control condition were also clearly observed in our gene expression analysis, as 2027 (960 up- and 1064 down-regulated) genes showed differential response between both the genotypes (Figure 1A). Further, GRN revealed calcium-binding protein CML27, ERF9, MYB1, NDR1/HIN113, cysteine-rich and transmembrane domain-containing protein B as highly connected hub genes, potentially regulating the basal defence mechanisms (Figure 6A). The relatively high expression of these genes compared to susceptible genotype indicates the pre-activated defence in AB-resistant chickpea genotype.

### Cell wall thickening and secondary metabolite biosynthesis mitigate infection at early stage

In lightcyan1 module, hub genes were mainly associated with cutin, suberin, wax and lignan biosynthesis. These genes encode for fatty alcohol:caffeoyl-CoA acyltransferase, 3-ketoacyl-CoA synthase, alkane hydroxylase MAH1, secoisolariciresinol dehydrogenase, and diacylglycerol O-acyltransferase. Besides, pleiotropic drug resistance protein 1 (PDR1), tyrosine decarboxylase1, and PRE6 were also found highly connected in the network (Figure 6B). PRE6 is an important transcriptional repressor of auxin response factors (Zheng et al., 2017). Tyrosine decarboxylase is involved in catecholamines biosynthesis (Świędrych et al., 2004). PDR1 participates in both constitutive and JA-induced defence against *B. cinerea* (Stukkens et al., 2005). Moreover, Cytochrome P450 oxidoreductase CYP736A12 and BRO1-domain containing protein were also present in this module. The BRO1-domain containing protein is induced in ER-stressed rice seeds (Qian et al., 2015). Although, no significant module was identified in susceptible genotype at 24-hpi, genes related with toxin, phytoalexins and secondary metabolite biosynthesis were over-represented among DEGs (Dataset S2). These data suggest that faster activation of cell wall thickening and defence-related secondary metabolite biosynthesis is crucial for early host response, and thus providing first line of defence in chickpea against *A. rabiei*.

### Signaling, protein phosphorylation and transcriptional machinery is majorly involved in coordinating AB resistance

At later stage (72-hpi), majorly PKs (32) and TFs (44) were over-represented. Resistant genotype-associated modules harbor 28 up-regulated RLK-Pelle members including LRR, DLSV and LRK, while, 12 down-regulated receptors mainly LRRs (Dataset S9). Mitogen-activated protein kinases (MAPK4 and MAPK11) were also induced, while MAPK9 was down-regulated. This indicates extensive involvement of post-translational regulation via protein phosphorylation at current or later stages. WRKY, NAC, bZIP and MYB TFs which are known to play important role in plant immunity were also induced along with other TFs *viz.* B3, GARP-G2-like, MADS-MIKC whose functions in defence remain elusive. Down-regulation of AP2/ERF, C2C2-Dof, and LIM TFs suggests that they could be negative regulators of AB-stress. Further, GRN identified plastid movement impaired protein, NAD(P)H-dependent 6’-deoxychalcone synthase, MYB14, chitinase and vestitone reductase hub genes suggesting that these could be master regulators of overall chickpea response to *A. rabiei* (Figure 6C).

Intriguingly, OPR11, SAMT, AdoMet synthase and SDR that are involved in phytohormone pathways were commonly identified between hub and DEGs. PR (PR-1B, PR-4, PR-5a, PR-10), E3 ubiquitin protein ligase DRIP2, thaumatin-like protein, and GSTs along with calcium signaling-related genes were also activated. In accordance to 24-hpi, coumarate-CoA ligase, feruloyl-CoA orthohydorxylase and isoflavone-2’-hydroxylase, involved in biosynthesis of phenylpropanoids and phytoalexins (Akashi et al., 1998; Kai et al., 2008; Liu et al., 2014; Schmid et al., 2014), were also present among the hub genes of 72-hpi. Together, our findings suggest that hub genes might have major role during AB disease, and importantly, enrichment of defence-related genes in resistant genotype compared to susceptible explains the contrasting behavior of both genotypes during fungal colonization.

In skyblue network, protein trichome birefringence 36, peroxidase, probable fructokinase, tubulin β-chain and 2-oxoglutarate-dependent dioxegenase were identified as hub genes (Figure 6D). In addition, susceptibility-associated genes *viz.* pectate lyase, polygalacturanse1, PR1, and Avr9/Cf-9 were identified. Pectate lyase (PMR6) confers powdery mildew susceptibility to arabidopsis (Vogel et al., 2002), whereas, polygalacturanse1 suppresses the programmed cell death (He et al., 2019). Importantly, down-regulation of *SWEET* transporter in resistant genotype further substantiates its role in disease susceptibility, as observed in earlier report (Cohn et al., 2014). *ENHANCED PSEUDOMONAS SUSCEPTIBILTY1* (*EPS1*), which provides SA-dependent resistance (Zheng et al., 2009), is down-regulated as SA-mediated defence is suppressed during necrotrophic infections. Intriguingly, development-related hub genes such as *FANTASTIC4*, *B-like cyclin*, *FT-interacting protein1* were also identified in down-regulated module implying growth-immunity trade-off.

### Regulators coordinating AB-susceptibility response in chickpea

The hub genes identified in GRN of susceptible genotype (darkmagenta) were-fasciclin-like arabinogalactan protein12, UTP-glucose-1-phosphate uridylyltransferase, cellulose synthase, DUF1336 family protein, and putative aminoacyltransferase (Figure 6E). Importantly, role of these genes is underexplored in plant defence/susceptibility, thus, they could be novel determinants for AB disease. PKs such as CAMK_CAMKL-CHK1/2, RLK-Pelle_LRR-XI-1/2, and bZIP, NAC, NF-YB TFs were identified among the hub genes. In addition, serine/threonine kinases, peroxidase, polyphenol oxidase A1, COBRA, putative senescence regulator S40 also responded to *A. rabiei*. The polyphenol oxidase A1 regulates secondary metabolism and cell death in walnut (Araji et al., 2014). Loss-of-function of COBRA invokes JA-regulated defence in arabidopsis (Ko et al., 2006). However, putative senescence regulator S40 is activated in response to natural senescence, darkness, pathogen, SA and ABA treatment (Krupinska et al., 2002). Overall, this module contains novel susceptibility factors and also suggests minimal activation of immunity in susceptible genotype, which was unable to mitigate the disease.

## DISCUSSION

To combat invading pathogen, plants modulate their growth, development and defence via complex and dynamic transcriptional alterations which start with perception and last till the defeat of pathogen or host (i.e. resistance or disease). Hence, the initial stages of infection are critical for the establishment of pathogen inside host and disease development (Mine et al., 2018). In chickpea, the early defence signaling and regulatory networks differentially operating in resistant and susceptible cultivars upon AB challenge remain elusive. Here, we investigated the pathogen-induced host transcriptional reprogramming occurring at early stages of *A. rabiei* infection in well-characterized AB resistant (FLIP84-92C) and susceptible (PI359075) chickpea genotypes using RNA-seq approach. Our comprehensive analysis provides an overview of chickpea defence response to a fungal necrotroph (Figure 7), and identifies differentially regulated genes and co-expression modules containing receptors, PKs, TFs along with other biosynthetic enzymes. In addition, novel hub genes were defined that are potentially responsible for the resistance/susceptibility of chickpea genotypes under AB stress.

**Figure 7:**
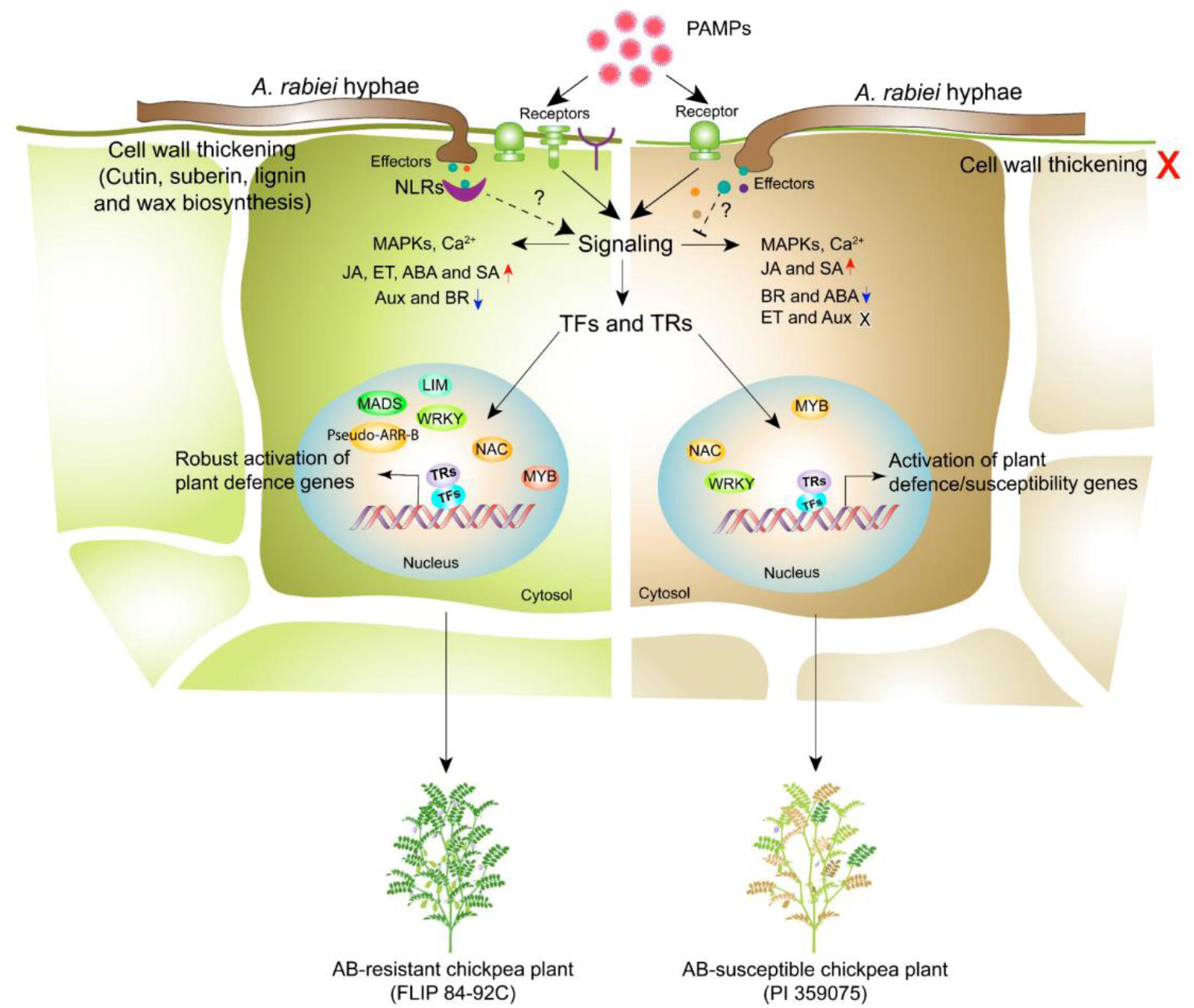
Early-stage transcriptional response of resistant and susceptible chickpea plants against *A. rabiei*. Schematic model highlighting the summary of cascades incurred during the differential interaction of *A. rabiei* with resistant (FLIP 84-92C; left) and susceptible (PI 359075; right) chickpea genotypes. Resistant plants deploy large number of receptor kinases to detect the pathogen-associated molecular patterns (PAMPs) compared to susceptible genotype. Besides, resistant plants activate genes controlling cell wall thickening process to limit the pathogen invasion. In contrast, the expression of cell wall thickening genes remain unaffected in susceptible genotype. Further, to overcome the host defence response pathogens utilise secreted effector proteins to suppress host-immune machinery in susceptible plants. However, resistant plants produce NLRs that recognise these effector molecules and activate downstream signaling. These defence signaling cascades involve a number of protein kinases, phytohormones and transcription factors/regulators (TFs/TRs). In resistant plants JA, ABA, and ET biosynthesis and regulatory pathways were induced, however, susceptible genotype only activated JA regulatory components. SA showed up-regulation in both the genotypes. The Aux and BR signaling pathways are down-regulated in resistant genotype. In comparison to susceptible plants, a vast variety of TFs and TRs such as WRKY, MYB and MADS-box were identified in resistant plants, which mediate downstream transcriptional activation of robust and complex defence mechanisms for successful neutralization of *A. rabiei*. On the contrary, susceptible plants could activate very less defence-related genes along with induction of susceptibility factors, leading to successful establishment AB disease.

A plethora of proteins participate in basal and induced processes of multi-layered immune system. The pre-formed or constitutively elevated expression of plant defence genes plays a major role in preventing pathogen invasion (Vergne et al., 2010). The wide-spectrum differential gene regulation between AB resistant and susceptible chickpea genotypes under control condition, suggests the existence of a functional and efficient pre-formed immune system in resistant genotype. However, the pre-activated basal defence components in susceptible genotype are not enough to restrict fungal invasion, and might serve as susceptibility factors. Extensive cell wall reinforcement and production of anti-microbial compounds are non-specific plant responses which act as first barrier against pathogen (Muthappa et al., 2013). They prevent or delay fungal penetration, thus, buy time for the activation of downstream defence system. At 24-hpi, mainly lignan, cutin, wax, suberin, and secondary metabolite biosynthesis-related genes were significantly induced, resulting in cell wall thickening. This data complements the earlier histological observations of physical barriers and cuticle layer in AB resistant chickpea cultivars (Ilarslan and Dolar, 2002). Intriguingly, high-amplitude transcriptional reprogramming in resistant genotype, especially at 72-hpi, highlighted the importance of this time-point in deciding the fate of *A. rabiei* colonisation. However, data for later stages would be vital to pin-point the minimum duration of such a high transcriptional reprogramming. Both genotypes shared small sub-set of DEGs at 24- and 72-hpi, that suggests the differential response of these cultivars against AB-stress. Enrichment analysis demonstrated extensive nexus of signaling pathways, post-translational modifications, phytohormone cross-talk, and secondary metabolism in resistant genotype to overcome pathogen.

Plant’s surveillance system against pathogens mainly relies on RLKs and NOD-like receptors (NLR) (Wang et al., 2020; Tang et al., 2017). Compared to susceptible genotype, high number of RLK-pelle members were identified in resistant genotype upon AB stress, whereas, none of them was induced in susceptible genotype at 24-hpi. Moreover, rust resistance kinase Lr10, receptor-like protein EIX1, receptor-like protein kinase HSL1 and wall-associated receptor kinase 20 were identified among hub genes under resistant modules. Lr10 is a CC-NBS-LRR receptor involved in immunity against wheat rust disease (Feuillet et al., 2003). MAPK signaling cascades triggered upon pathogen recognition act on WRKY TFs which are regulators of host defence responses against a variety of pathogens (Eulgem and Somssich, 2007). In our datasets also several MAPKs and WRKY TFs, especially WRKY4, WRKY47, WRKY31 and WRKY65 were mainly induced in FLIP84-92C. Besides, MADS-Box, MYB, bHLH and B3 TFs were also preferentially expressed in resistant genotype. TFs play crucial role in biotic stress response through complex GRNs (Park et al., 2001; Wang et al., 2009). In our study, MYB14 and PRE6 showed highest connectivity, hence plausibly modulating other genes of the resistant module. Furthermore, the abundance of SOC1 and BPC1 targets such as disease resistance protein RPS5, PR-genes and other defence-related transcripts among the DEGs, prospects them as master regulators of chickpea immunity against *A. rabiei*, apart from their developmental functions. Notably, PR-1B, PR4, PR-5a and PR-10 were also identified as hub genes in resistant plants. PR-10 provides defence by preventing/disrupting the conidial germination and hyphal growth (Zandvakili et al., 2017). In arabidopsis enhanced disease susceptibility (*eds*) mutants, the altered PR1 expression enhances susceptibility to bacterial pathogens (Rogers and Ausubel, 1997). Together, results concluded that resistant cultivar triggered prompt and robust defence via reprogrammed expression of signaling and transcriptional regulatory components in response to *A. rabiei*. On the contrary, susceptible genotype could not mount sufficient response to mitigate the disease progression. The minimal activation or suppressed immunity could be attributed to the secreted effector proteins as *Ascochyta* genome encodes an armory of predicted effector proteins (Verma et al., 2016b).

SA and JA are classically known to regulate signaling networks involved in induced defence responses against biotrophic and necrotrophic pathogens, respectively (Ryan and Moura, 2002; van Loon et al., 2006; Abd El Rahman et al., 2012). In response to *A. rabiei,* extensive modulation of phytohormone pathways occurred in host. Our data demonstrates induced expression of JA biosynthesis genes in resistant genotype; however, SA biosynthesis remained unchanged. Importantly, ET and ABA biosynthesis pathways were also induced which are not well known for defence against necrotrophic fungi. This indicates the possibility of their interaction with JA signaling to combat AB stress. The physiologically active hormone levels are controlled through fine-tuning of *de novo* biosynthesis and catabolism (Saito et al., 2004). Intriguingly, differential regulation of hormone conversion enzymes such as ABA hydroxylase, JMT, SAMT and methylxanthosine synthase 1 occurred upon *A. rabiei* challenge. SAMT converts SA into MeSA and methylxanthosine synthase 1 plays crucial role in alkaloid biosynthesis (Mizuno et al., 2003; Deng et al., 2020). This suggests active conversion of SA into other forms during infection. Higher expression of JMT, which converts JA into MeJA, mediates jasmonate-regulated plant defence against a necrotrophic fungus *Botrytis cinerea* (Seo et al., 2001). Nowadays, auxin is considered as stress-responsive hormone besides its role in plant growth and developmental (Wang and Fu, 2011). The down-regulation of auxin-related genes suggests suppression of growth response while promoting defence, thus indicating antagonistic nature of auxins with defence hormones. Together, these hormonal modulations cumulatively orchestrate the early defence response in chickpea and confer AB resistance.

Furthermore, genes involved in plant growth, flowering time, and development were also identified among the DEGs and hub genes. Interestingly, chloroplast and photosynthesis-related genes showed up-regulation in resistant genotype, however, remain unchanged in susceptible. This could be possibly mediated by fungal effectors as many effector proteins localize to chloroplast, and interfere with photosynthetic processes or ROS levels by interacting with target proteins, thus, promoting susceptibility (Singh et al., 2021b). Moreover, leaf and floral development genes have been located on the QTL regions conferring AB resistance (Daba et al., 2016; Kumar et al., 2018; Deokar et al. 2019). This emphasises on the balanced resource allocation between pathogen combat machinery and growth regulation. Most likely these genes are employed in the well-known phenomenon of growth-immunity trade-off. However, their functional correlation with chickpea immunity and development needs to be investigated. It will also be interesting to find how pathogen infection affects floral development or *vice-versa*.

In summary, we report the comprehensive transcriptional landscape during *Cicer-Ascochyta* interaction, and divulge the key mechanisms underlying AB resistance/susceptibility in contrasting chickpea genotypes. In addition, we identify early immune signaling components and highlight possible candidate genes that may play important role in disease resistance. Although, the molecular function of identified hub genes requires further experimental investigations, our data strongly indicates their critical role in orchestrating chickpea immunity. Taken together, this study improves our understanding of AB disease development and delivers a handful of candidate genes to be exploited in the genetic manipulation or breeding programs for chickpea improvement.

## Author contributions

PKV and RS conceptualise and designed the experiments. RS performed the wet lab experiments. RS and AD analysed the RNA-seq data. RS, YS and KK interpreted the data and wrote the draft manuscript. AR and PKV critically revised the manuscript. PKV arranged the funds.

## Data availability

All relevant data can be found within the manuscript and its supplementary materials. Please contact the corresponding author to obtain raw data from RNA-seq analysis.

